# Dual-Coil Transcranial Magnetic Stimulation Reveals Temporal Dynamics of Bilateral Corticomotor Excitability During Response Inhibition

**DOI:** 10.1101/2022.09.14.507942

**Authors:** Chantelle E. Ratcliffe, Craig J. McAllister, Hayley J. MacDonald

## Abstract

A changing environment may suddenly require some parts of a multi-component response to be cancelled, while others continue. Such partial cancellation consistently produces a behavioural delay in the remaining component. This delay may reflect a three-step process of non-selective neural inhibition of all response components, functional uncoupling of components, and selective initiation of the remaining response. However, most neurophysiological evidence supporting this hypothesis has been recorded from muscles of a single hand, without direct comparison between response components. We aimed to simultaneously record - and therefore directly compare - corticomotor excitability (CME) in the cancelled and responding hands using a dual-coil technique not yet applied in this context. Human participants received transcranial magnetic stimulation to both primary motor cortices 1ms apart while performing a bimanual response inhibition task. Motor evoked potentials (MEPs) were recorded from both first dorsal interosseous (FDI) muscles as a measure of CME during partial cancellation. An equivalent reduction in CME was evident for both the responding and cancelled FDI muscles 175 ms after the stop cue during successful partial cancellation. The responding FDI subsequently exhibited an increase in CME above levels in the cancelled hand, leading to the unimanual response. This study reveals, for the first time, the temporal dynamics of CME for both response components simultaneously during a bimanual response inhibition task. Our results provide strong evidence that neural modulation during sudden partial cancellation can be viewed in light of the stop-change framework as a sequential non-selective stop, uncouple and switch, then selectively go process.

## Introduction

Flexibly adapting behaviours in an ever-changing environment is a core aspect of cognitive control in everyday life. Some situations require the sudden and complete cancellation of prepared action i.e. complete response inhibition. More complex scenarios require only some parts of a multi-component action to be cancelled while others continue i.e. partial response inhibition. Partial response inhibition consistently produces a substantial behavioural delay in the continuing component, often termed a stopping interference effect e.g., (Aron & Verbruggen, 2008; Coxon, Stinear, & Byblow, 2007; Ko & Miller, 2011; Wadsley, Cirillo, & Byblow, 2019). However, the neural mechanisms contributing to this effect are still disputed.

Response inhibition engages right-lateralised inhibitory control networks comprising cortical, basal ganglia, and thalamic regions (Aron & Poldrack, 2006; Rubia, Smith, Brammer, & Taylor, 2003; Zandbelt, Bloemendaal, Hoogendam, Kahn, & Vink, 2013). These subcortical networks ultimately decrease excitability within the primary motor cortex (M1) (Zandbelt et al., 2013) to suppress descending output to task-relevant muscles and cancel planned movements. Transcranial magnetic stimulation (TMS) is commonly used to study task-dependent effects on corticomotor excitability (CME), as measured via amplitude changes in the motor evoked potentials (MEPs) of target muscles, during impulse control tasks. Traditionally, during any single trial of a response inhibition task, CME is recorded from muscles on one side of the body using one TMS coil. Therefore, any bilateral between-muscle comparisons are made from MEPs collected on separate trials. The relatively large trial to trial variability in MEP amplitude can make this problematic. However, a dual-coil TMS technique has recently been developed to record CME bilaterally, stimulating both M1s with a 1 ms inter-pulse interval (Grandjean et al., 2018; Wilhelm, Quoilin, Petitjean, & Duque, 2016). The 1ms interval avoids direct electromagnetic interference between coils (Wilhelm et al., 2016) and does not activate any interhemispheric mechanisms that can occur with longer intervals (>4 ms) (Ferbert et al., 1992). This technique has successfully revealed CME dynamics for both hands simultaneously during movement preparation in a choice reaction time task (Vassiliadis et al., 2018). To our knowledge, we are the first study to use this technique to directly compare bilateral CME dynamics within the same trial during a response inhibition task.

Mounting evidence indicates that partial movement cancellation has a widespread effect on M1 beyond that of the cancelled components; also influencing those representations corresponding to task-irrelevant (Majid, Cai, George, Verbruggen, & Aron, 2012) and homologous muscles (Ko & Miller, 2011). Of most relevance for the current study, output from M1 is suppressed 175 ms after the (irrelevant) stop cue for muscles of the remaining movement (Cowie, MacDonald, Cirillo, & Byblow, 2016; MacDonald, Coxon, Stinear, & Byblow, 2014). We have previously interpreted this neural suppression in a muscle that was never cued to stop as evidence of neuroanatomical constraints that limit the ability to selectively inhibit actions in this context i.e., when a functionally coupled bimanual response has been anticipated. Consequently, inhibition would have to be equally applied to all components of the initiated bimanual response. Therefore, confirmation that the observed suppression indeed reflects a non-selective inhibitory process necessitates suppression being observed concurrently, and at an equivalent magnitude, in both the responding and cancelled muscles. The most reliable way to test for this concurrent suppression is to record CME simultaneously from bilateral muscles.

The present study had two aims: 1) to validate the dual-coil TMS technique in the context of response inhibition, and 2) to use this technique to directly test the working hypothesis that partial response inhibition is associated with a non-selective stopping mechanism, by revealing CME dynamics of both the cancelled and responding components within the same trial. As an initial reproducibility test to address the first aim, we attempted to replicate the results of MacDonald et al. (2014) demonstrating that when the left finger responded during a partial cancellation trial, MEP amplitude decreased 175ms after the (irrelevant) stop cue, followed by an increase in amplitude leading to the behavioural response. The novel application of the dual-coil technique allowed us to test our new hypothesis that CME suppression would occur at the equivalent time point, 175ms after the stop cue, and to an equivalent degree in the right finger which was cued to stop. We further hypothesised that, following the simultaneous suppression, excitability for the left and right finger would diverge as CME would not need to increase for a right finger response. Finally, we hypothesised that during unsuccessful partial cancellation trials when a bimanual response was incorrectly produced, CME would be higher in both fingers and therefore not suppressed to the same level compared to successful partial cancellations trials.

## Materials and Methods

### Participants

Twenty-three healthy adults (mean age 20.3 years, range 18 - 35, 19 female) with no neurological impairment were recruited into this experiment. This recruitment target exceeded the sample size required (N = 15) for 0.8 power (Faul, Erdfelder, Lang, & Buchner, 2007) given the medium effect size (Cohen’s d = 0.69) relating to the MEP amplitude decrease 175ms after the stop cue during partial cancellation (MacDonald et al., 2014). All participants were right handed (laterality quotient mean 94, range 42 - 100) as assessed using the Edinburgh Handedness Inventory (Oldfield, 1971). All participants provided written informed consent and completed a TMS screening questionnaire based on current safety guidelines (Rossi, Hallett, Rossini, Pascual-Leone, & Safety of, 2009). Ethical approval was obtained from the University of Birmingham Ethics Committee (ERN_17-1541AP1).

### Experimental Design

#### Behavioural task

Partial response inhibition was investigated using the bimanual anticipatory response inhibition (ARI) task first described by Coxon et al (2007). Participants were seated in front of a monitor (48 × 27 cm) displaying two vertical white rectangles (Figure 1A). Two custom-made switches controlled two rising black bars within these rectangles; the left bar corresponded to the left switch and the right bar to the right switch. Participants were instructed to depress both switches with the weight of their index fingers to commence each trial. After a random delay, ranging from 400 – 900 ms, both bars would rise vertically at a constant speed, reaching a horizontal target line at 800ms, and the top of the rectangle at 1000ms. Participants were informed that the bar(s) would cease rising when the corresponding index finger was lifted from the switch. The lift response required index finger abduction via contraction of the first dorsal interosseous (FDI) muscle. This was detected via the state of the custom-made microswitches monitored through an Arduino device and synchronised to the visual display via an analogue-to-digital USB interface (NI-DAQmx 9.7; National Instruments).

**Figure 1.**
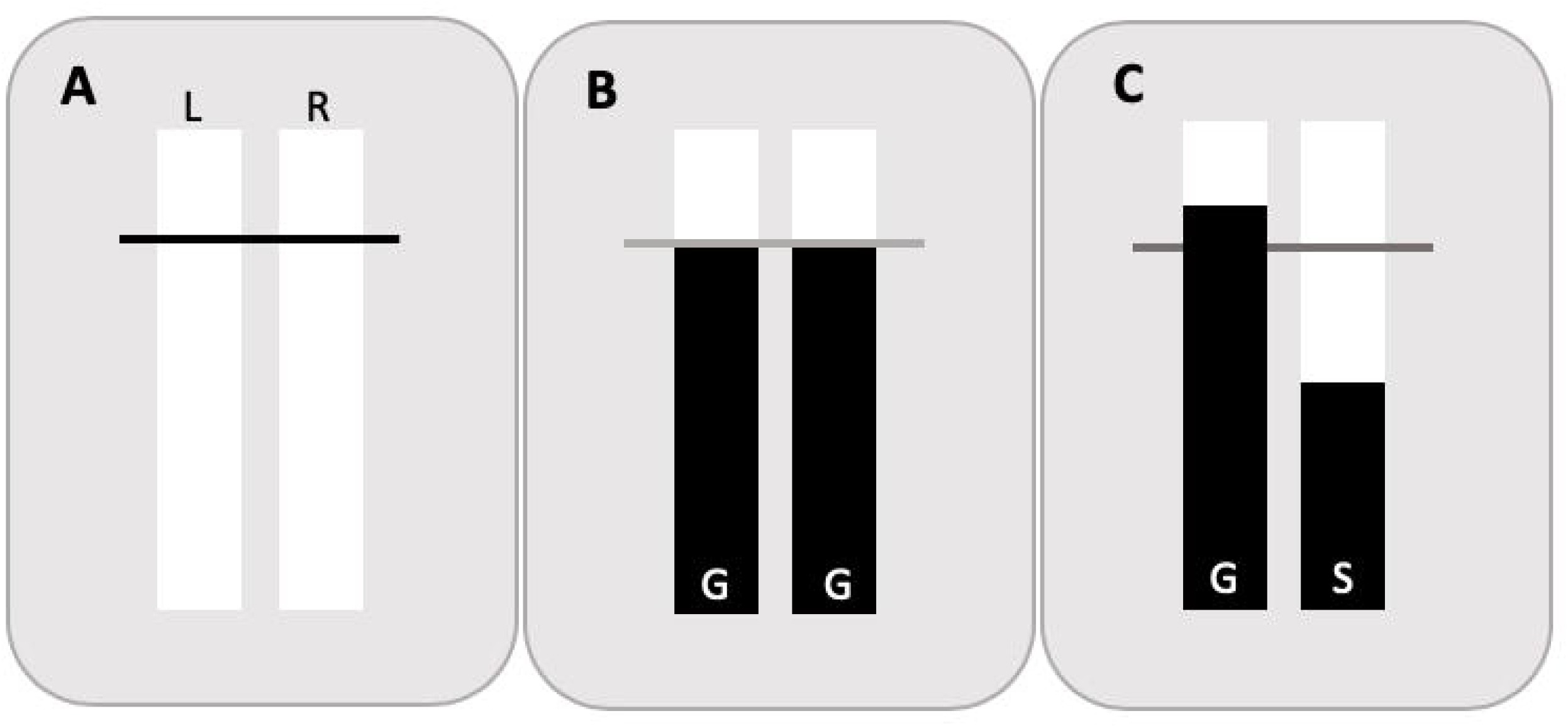
Behavioural task. A) Visual display shown at the start of the trial - trial type is unknown. Left (L) bar corresponds to left finger, right (R) bar corresponds to right finger. B) Successful Go (Go Left – Go Right: GG) trial – both fingers lifted to intercept both bars with the target line. C) Successful Go Left – Stop Right (GS) trial-right finger response was correctly inhibited whilst left finger response was lifted, but target is missed as lift response is delayed.

The default response (66% of trials) required participants to lift both index fingers from the switches to intercept both rising bars with the target line (Go Left – Go Right: GG trial; Figure 1B). The other type of trial was a Stop trial (33% of trials) in which the bars could automatically stop rising before reaching the target, which required the participant to inhibit their planned lift response. There were three types of Stop trials with set stop times. When both bars stopped automatically 200ms before the target, the participant was required to keep both fingers on the switches until the end of the trial (Stop Left – Stop Right: SS trial). When only one bar stopped rising 250ms before the target whilst the other bar continued to rise (partial response inhibition trials), the participant was required to hold the corresponding finger on the switch until the end of the trial whilst lifting the other finger from the switch to intercept the bar with the target line. Either the left (Stop Left – Go Right: SG trial) or right bar could stop early (Go Left – Stop Right: GS; Figure 1C). Bar stop times were chosen to allow direct comparison to previous findings (Cowie et al., 2016; MacDonald et al., 2014) and reflect the increased difficulty of partial compared to complete response inhibition, necessitating an earlier stop time. After each trial, visual feedback displayed whether the participant had responded correctly, and if the rising bar(s) had been stopped within 30ms of the target line. Trial order and visual output was controlled by custom software written in MATLAB (R2016a, version 9.0; The MathWorks).

All participants completed practice blocks, consisting of 35 GG trials only, before the main experimental blocks to ensure consistent accuracy for the default movement. The main task consisted of 432 trials, spread across 12 blocks of 36 trials, of which 288 were GG trials. 114 Go Left - Stop Right (GS) trials were the main trials of interest for direct replication and extension of previous work (Cowie et al., 2016; MacDonald et al., 2014). 15 Stop Left – Stop Right (SS) trials and 15 Stop Left - Go Right (SG) trials were included as catch trials to ensure that participants were not able to predict a GS response. Due to the required number of additional trials (at least 198 SS and SS trials, plus 400 GG trials to maintain a 2:1 Go/Stop ratio), it was not feasible to measure equivalent CME curves in SS and SG trials within a single session for a direct comparison.

#### Electromyography

Surface electromyography (EMG) recordings were obtained from bilateral FDI muscles using a Delsys Bagnoli-2 EMG System, with the ground electrode placed on the elbow. EMG activity was amplified (x1000), band-pass filtered (20-450Hz) before sampling at 2000ms, using a Micro 1401 analogue to digital convertor (Cambridge Electronic Design, Cambridge, UK). The data was analysed on a PC using Signal version 6.04 software (CED). EMG was triggered by the Arduino and recorded for 1000ms once the bars began to rise in the ARI task.

#### Transcranial magnetic stimulation

Single TMS pulses were delivered through two separate 70mm figure-of-eight shaped coils that were connected to two separate Magstim 200^2^ stimulators (Magstim Company Ltd, Whitland, UK) and triggered with precise timing intervals via the Arduino device. Coils were placed over the right and left M1 with the handles pointing backward and laterally away from the midline at a 45-degree angle to produce current in an anterior-to-posterior direction. For each M1, the optimal coil position that produced the largest and most consistent motor evoked potentials (MEPs) in the relaxed contralateral FDI muscle was marked on the scalp to provide a reference throughout the experiment. TMS intensity was then gradually decreased to determine the motor threshold, which was defined as the minimal intensity that evoked MEPs of above 50μv peak-to-peak in 5 out of 10 consecutive trials. The test stimulus (TS) intensity for each FDI used throughout the experiment was determined during practice blocks, which consisted of only GG trials. TS was initially set at the participant’s motor threshold and adjusted in increments of 1 – 2 % maximum stimulator output as necessary to obtain MEPs of 200 – 400 μv when stimulation occurred 200ms before the target i.e., when the muscle was still quiescent (MacDonald et al., 2014). This relatively low stimulation intensity is less likely to disturb behavioural performance and is more sensitive to fluctuations in CME compared to higher intensities, due to predominantly indirect wave recruitment (Di Lazzaro, Oliviero, et al., 1998; Di Lazzaro, Restuccia, et al., 1998). Adjustment of TS was performed separately for each target muscle and the TS was then kept constant throughout the main experiment. During the experimental blocks of the ARI task, MEPs were elicited from both the left and right FDI with a 1 ms inter-pulse interval.

TMS was applied in 84 of the 288 GG trials at seven time-points (250, 225, 200, 175, 150, 125, or 100ms) before the target to capture modulation of CME in the lead up to movement execution. TMS was applied in 84 of the 114 GS trials at one of seven time-points (150, 125, 100, 75, 50, 25, or 0ms) before the target. Stimulation times were offset by 100 ms in GS trials as responses are robustly delayed on these trials compared to GG trials. Twelve MEPs were obtained from both FDI muscles at each time-point. No stimulation was applied in SS and SG catch trials.

#### Data analysis

All lift times were calculated in milliseconds relative to the target. Mean lift times were calculated for all GG trials, successful GS trials and unsuccessful GS trials after the removal of outliers in each condition (± 3 standard deviations). Lift times from GS trials correspond to the responding left finger. A GS trial was considered successful if the right finger remained down on the switch until the end of the trial, whilst the left finger lifted before the end of the trial. A GS trial was considered unsuccessful and included in the analysis if both fingers were lifted from the switches, mimicking a GG trial. Unsuccessful trials where both fingers remained on the switches or only the incorrect right finger was lifted from the switch, were not included in the analysis. Percentage of successful trials and stop signal reaction time (SSRT) were calculated for GS trials. SSRT was calculated using the integration method (Verbruggen, Chambers, & Logan, 2013) as stop times were pre-set rather than determined via the staircase algorithm.

Baseline EMG activity was calculated as the root-mean-square amplitude in the 50 ms prior to the earliest TMS pulse. MEPs were excluded if this baseline EMG was above 15μV. MEPs were measured from both FDI muscles as the peak-peak amplitude in the 15-45ms after the TMS pulse. During GG and GS trials, mean MEP amplitude was calculated for each stimulation time-point after trimming upper and lower 10% of MEPs (if > 10) for a more accurate measure of centrality (Wilcox, 2001). Given the possibility of lower MEP numbers at some stimulation times, the median MEP amplitude was also calculated for comparison to ensure mean data was not skewed.

#### Statistical analysis

One-way repeated measures analysis of variance (RM-ANOVA) with post hoc comparisons compared the lift times for the left finger during GG, successful GS and unsuccessful GS trials.

RM-ANOVAs were also used to test the hypotheses relating to the CME dynamics and baseline EMG. A 2 Side (Left, Right) × 7 Stimulation Time (−250, −225, −200, −175, −150, −125, −100ms relative to target) RM-ANOVA tested for differences in GG trials. On successful GS trials, MEP amplitudes and baseline EMG were analysed with a similar 2 Side × 7 Stimulation Time (−150, −125, −100, −75, −50, −25, −0ms relative to target) RM-ANOVA to compare CME dynamics between fingers. A planned comparison in MEP amplitude on GS trials between timepoints −100 ms and −75 ms tested for the CME suppression 175 ms post stop signal observed previously (Cowie et al., 2016; MacDonald et al., 2014). Post-hoc t-tests were performed with Sidak correction for multiple comparisons where necessary.

Differences in MEP amplitude and baseline EMG between successful and unsuccessful GS trials were investigated with a three-way RM-ANOVA with a 2 Success (Successful, Unsuccessful) × 2 Side × 7 Stimulation Time design.

Values are reported as means ± standard error (SE). Statistical significance was determined by α ≤ 0.05. Data which violated the assumption of sphericity are reported with Greenhouse-Geisser *P* values. Both partial and generalized eta squared effect sizes are reported for significant and approaching significant (0.1 > *P* > 0.05) ANOVA results, to allow future comparisons to both similar and different experimental designs (Lakens, 2013). Generalized eta squared effect sizes can be interpreted according to traditional benchmarks (small = 0.01; medium = 0.06; large = 0.14) (Cohen, 1988; Lakens, 2013).

## Results

The pre-set stop time of 250ms prior to the target on GS trials replicated previous methodology to enable direct comparisons (Cowie et al., 2016; MacDonald et al., 2014). However, five participants in the current study were unable to complete enough successful GS trials at the pre-set stop time to provide MEPs at every stimulation time. Behavioural and neurophysiological results are therefore reported for the remaining 18 participants unless otherwise stated. A sample size of 18 was still more than sufficient to achieve 0.8 power (Faul et al., 2007).

### Behavioural data

Mean lift times recorded for the left finger during GG, successful GS and unsuccessful GS trials are presented in Table 1. There was a main effect of Trial Type on lift times of the left finger (*F*_2,34_ = 56.990, *P* < 0.001, □_p_^2^ = 0.770, □_G_^2^ = 0.52). Post-hoc comparisons revealed that the lift time during successful GS trials (71 ± 6ms) was delayed compared to GG trials (14 ± 4ms, *P* < 0.001). Participants therefore showed the robust lift time delay on successful GS trials compared to GG trials. Lift times from successful GS trials were also delayed compared to the left finger on unsuccessful GS trials when a bimanual response was made in error (34 ± 6ms, *P* < 0.001). Finally, the left lift time from unsuccessful GS trials was delayed relative to the default GG trials (*P* = 0.040), even though the behaviour observed was the same (i.e. a bimanual lift response).

**Table 1.** Behavioural lift times for the left digit across trial types. Lift times (ms) for the left finger during Go Left – Go Right (GG) trials, successful Go Left – Stop Right (GS) trials and unsuccessful GS trials, for each participant. Lift time is relative to target at 800ms.

| Participant | GG | Successful | Unsuccessful |
| --- | --- | --- | --- |
|  |  | GS | GS |
| 1 | 35 | 96 | 62 |
| 2 | 18 | 68 | 46 |
| 3 | 3 | 72 | 19 |
| 4 | -6 | 36 | 17 |
| 5 | -5 | 32 | 13 |
| 6 | 18 | 68 | 75 |
| 7 | 23 | 83 | 78 |
| 8 | 5 | 87 | 14 |
| 9 | 50 | 104 | 20 |
| 10 | 26 | 92 | 12 |
| 11 | 7 | 85 | 49 |
| 12 | -6 | 6 | -14 |
| 13 | 7 | 108 | 47 |
| 14 | 19 | 67 | 52 |
| 15 | 1 | 66 | 24 |
| 16 | 27 | 80 | 33 |
| 17 | -2 | 37 | 15 |
| 18 | 28 | 76 | 46 |
| Average | 14 | 71 | 34 |
| SD | 16 | 27 | 24 |
| SE | 4 | 6 | 6 |

Stopping performance on GS trials was as expected and replicated previous findings. The success rate at the pre-set stop time during GS trials was 57 ± 4% and the SSRT was 231 ± 8ms.

### Neurophysiological data

#### GG trials (N=18)

Figure 2A shows the mean MEP amplitude at each stimulation timepoint for the left and right FDI muscles during GG trials. There was a main effect of Stimulation Time (*F*_6,102_ = 8.660, *P* < 0.001, □_p_^2^ = 0.337, □_G_^2^ = 0.09). MEP amplitudes gradually increased in both FDI muscles as the target line approached and participants prepared to lift with both fingers. However, as evidenced by the lack of a Side effect (*F*_1,17_ = 0.356, *P* = 0.559) or Side × Stimulation Time interaction (*F*_6,102_ = 0.666, *P* = 0.595), the rise in CME did not differ between FDI muscles. Collapsed across side (Figure 2B), MEP amplitude first increased relative to −250ms baseline levels (0.201 ± 0.03mV) at −150ms (0.297 ± 0.04mV; t_17_ = −3.614, *P* = 0.044).

**Figure 2.**
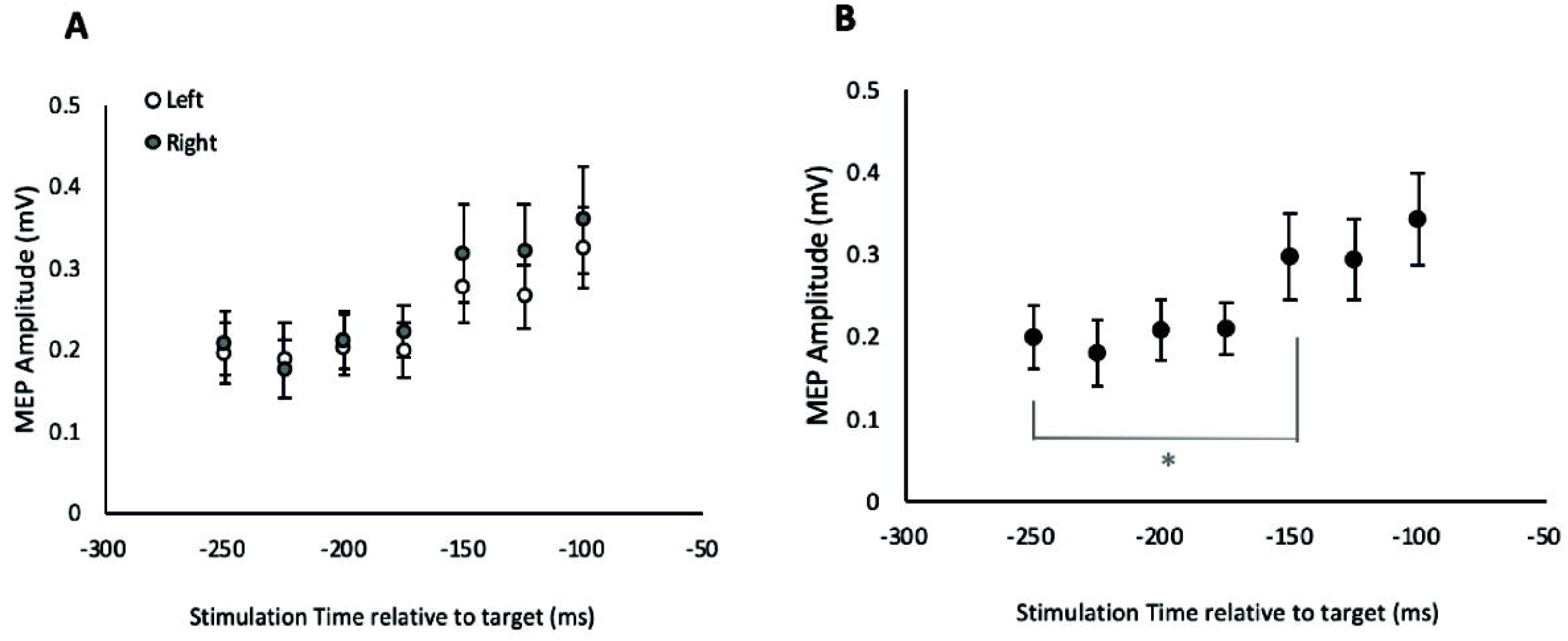
Modulation of corticomotor excitability during bimanual execution. A) Mean motor evoked potential (MEP) amplitude for left and right FDI during GG trials. B) Mean MEP amplitude collapsed across Side during GG trials. Values are means ± standard error. * represents significant difference (*P* < 0.05)

Median MEP amplitude showed the same pattern of results with a main effect of Stimulation Time (*F*_6,102_ = 6.968, *P* < 0.001, □_p_^2^ = 0.291, □_G_^2^ = 0.09), no main effect of Side (*F*_1,17_ = 0.109, *P* = 0.745) or Side × Stimulation Time interaction (*F*_6,102_ = 0.404, *P* = 0.802).

Main effects and interactions for baseline EMG were not statistically significant (all *P* > 0.1) so changes in background muscle activity cannot account for the changes in MEP amplitude. Baseline EMG collapsed across timepoints was 2.0 ± 0.04 μV for left and 1.9 ± 0.04 μV for right FDI muscles.

#### Successful GS trials (N=18)

Figure 3A shows mean MEP amplitude for the left and right FDI muscles at each stimulation timepoint during successful GS trials. There was a main effect of Stimulation Time (*F*_6,102_ = 5.G207, *P* = 0.007, □_p_^2^ = 0.234, □_G_^2^ = 0.07) and a planned comparison confirmed that MEP amplitude decreased from −100 ms (0.312 ± 0.06mV) to ±75ms (0.239 ± 0.05mV; t_17_ = 3.104, *P* = 0.006; Figure 3B) across muscles, including a replication of MEP amplitude suppression in the left FDI (0.338 ± 0.06mV to 0.265 ± 0.06mV; t_17_ = 2.522, *P* = 0.022). As the stop signal was presented 250ms prior to the target, the equivalent MEP decrease in both the responding (left) and cancelled (right) FDI muscle occurred 175ms after the stop signal was presented.

**Figure 3.**
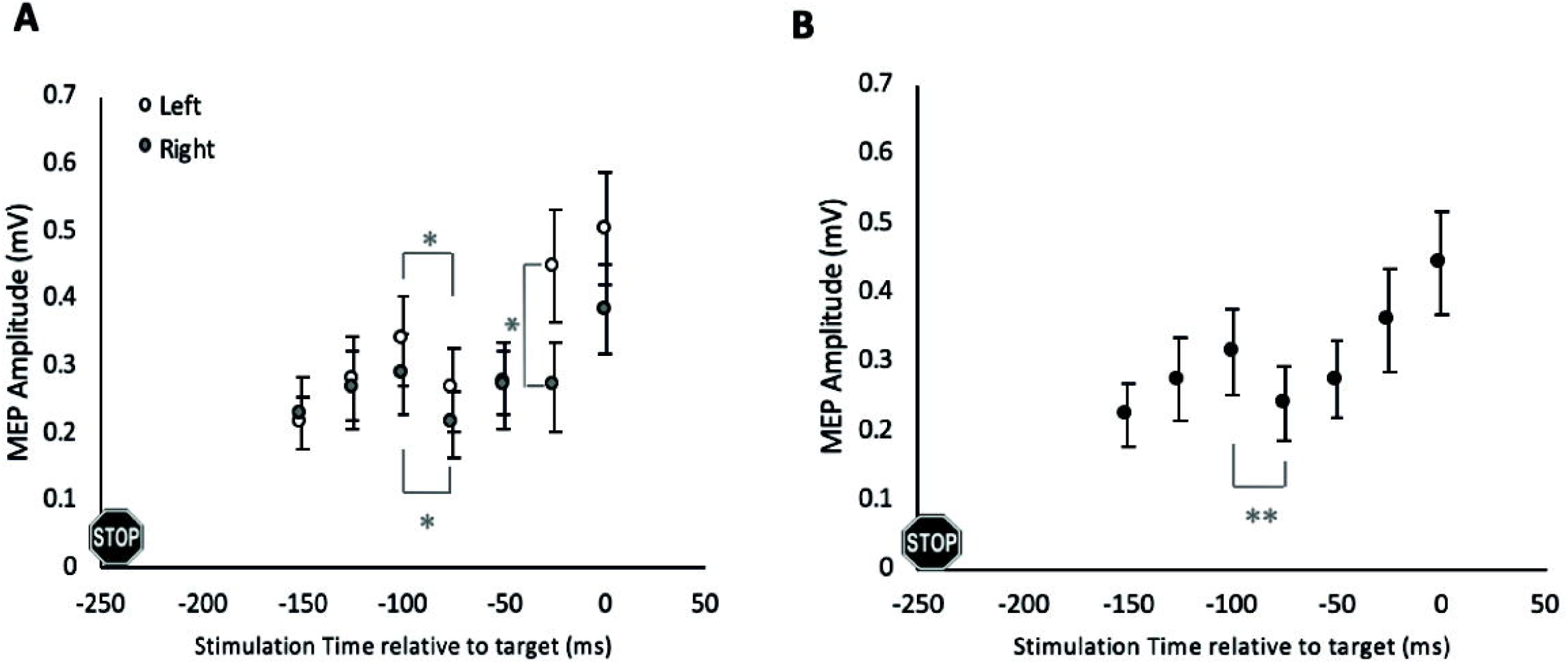
Modulation of corticomotor excitability during successful partial response inhibition. A) Mean motor evoked potential (MEP) amplitude for left (responding) and right (cancelled) FDI during successful GS trials. B) Mean MEP amplitude collapsed across Side during successful GS trials. Stop signal presented 250 ms before target. Values are means ± standard error. Significant differences during GS trials represented with * (*P* < 0.05) ** (*P* < 0.01)

There was no main effect of Side (*F*_1,17_ = 1.729, *P* = 0.206), but the Side × Stimulation Time interaction approached significance (*F*_6,102_ = 2.721, *P* = 0.059, □_p_^2^ = 0.139, □_G_^2^ = 0.02). Both the left (*P* = 0.022) and right (*P* = 0.040) FDI muscles showed the decrease in MEP amplitude at −75 ms, but then began to rise again. However, the trend towards a significant interaction arose as the increase in the responding left FDI was higher than the cancelled right FDI with a significant difference at 25 ms prior to the target (left: 0.448 ± 0.08mV, right: 0.270 ± 0.07mV; t _17_ = 2.557, *P* = 0.02; Figure 3A). Despite this divergence appearing to continue at 0ms, increased variability at this time point led to a non-significant difference (left: 0.502 ± 0.08mV, right: 0.383 ± 0.07mV; t _17_ = 1.781, *P* = 0.093).

Median MEP data supported these novel results by confirming a Side × Stimulation Time interaction (*F*_6,102_ = 3.844, *P* = 0.012, □_p_^2^ = 0.184, □_G_^2^ = 0.03), a main effect of Stimulation Time (*F*_6,102_ = 3.910, *P* = 0.011, □_p_^2^ = 0.187, □_G_^2^ = 0.06), and no effect of Side (*F*_1,17_ = 2.802, *P* = 0.112).

The changes in MEP amplitude were not driven by changes in background muscle activity. For mean baseline EMG, there was no effect of Side (*F*_1,17_ = 1.384, *P* = 0.256) but an effect of Stimulation Time (*F*_6,102_ = 2.951, *P* = 0.035, □_p_^2^ = 0.148, □_G_^2^ = 0.03). When collapsed across Side, there was a linear trend (*P* = 0.028) as baseline EMG increased incrementally when participants were preparing to respond at the target, but no stimulation time point rose significantly above −150 ms (all *P* > 0.093). Baseline EMG was 2.6 ± 0.1 μV for left and 2.2 ± 0.1 μV for right FDI muscles when collapsed across timepoint. Crucially, there was no Side × Stimulation Time interaction (*F*_6,102_ = 0.689, *P* = 0.557) that could account for the novel pattern of MEP results.

#### Unsuccessful vs Successful GS trials (N=13)

A GS trial was considered unsuccessful and included in the analysis if both fingers were lifted from the switches, mimicking a GG trial. Unsuccessful trials where both fingers remained on the switches or only the incorrect right finger was lifted from the switch, were not included in the analysis. Five participants did not produce a sufficient number of the required unsuccessful GS trials and were therefore excluded from this analysis. Data is presented, and comparisons made against successful GS trials, for the remaining 13 participants.

For mean MEP data there was a main effect of Success (*F*_1,12_ = 19.524, *P* < 0.001, □_p_^2^ = 0.619, □_G_^2^ = 0.06), with higher MEP amplitudes across stimulation timepoints on unsuccessful (0.444 ± 0.07 mV) compared to successful trials (0.296 ± 0.05 mV; Figure 4). There was also a main effect of Stimulation Time (*F*_6,72_ = 6.046, *P* = 0.012, □_p_^2^ = 0.335, □_G_^2^ = 0.09) and a Success × Stimulation Time interaction that trended towards significance (*F*_6,72_ = 2.059, *P* = 0.070, □_p_^2^ = 0.146, □_G_^2^ = 0.01). There was no main effect of Side (*F*_1,12_ = 2.213, *P* = 0.163) or any other interactions (all *P* > 0.207).

**Figure 4.**
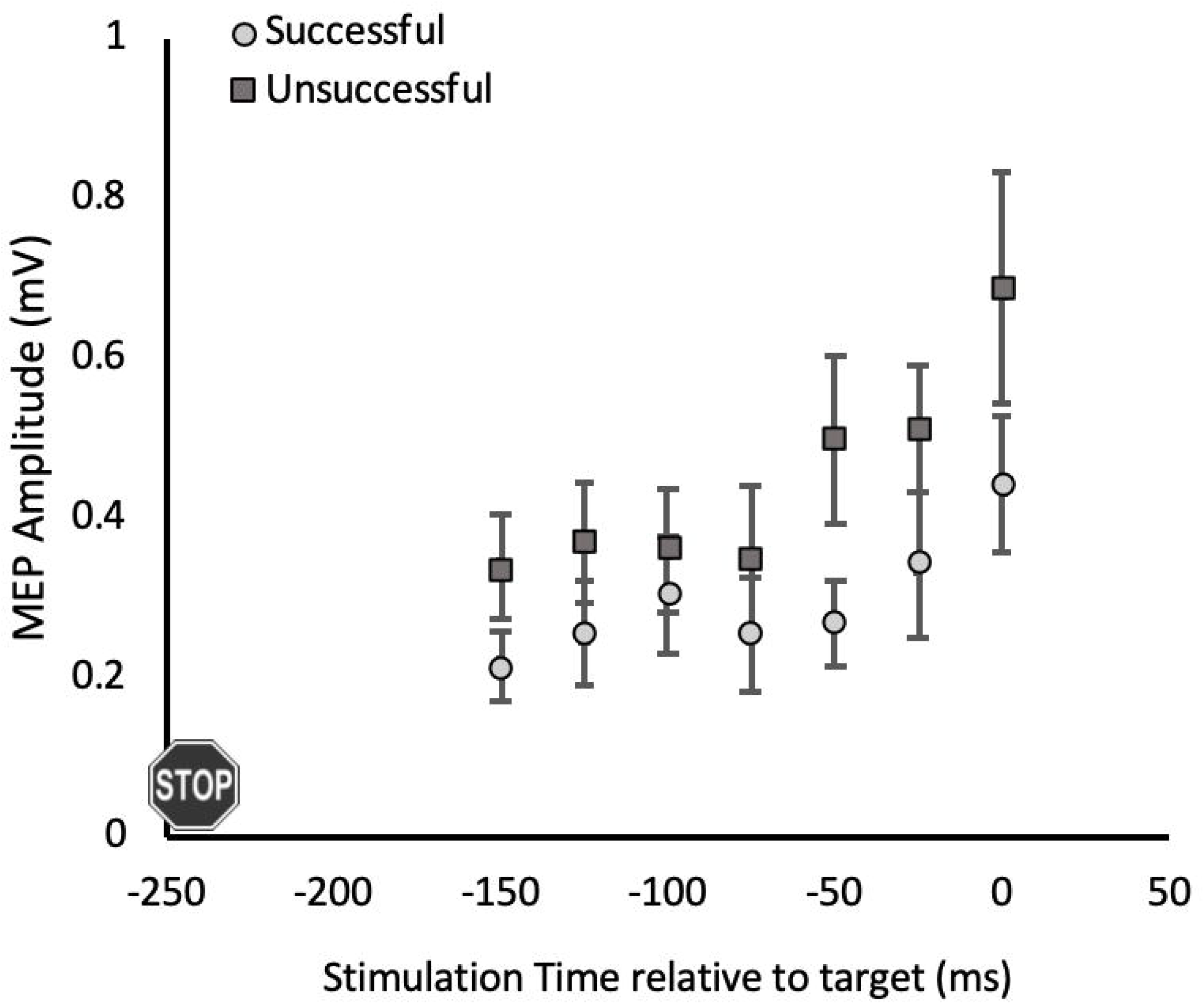
Comparison of corticomotor excitability during successful vs unsuccessful partial response inhibition. Mean motor evoked potential (MEP) amplitude for each stimulation timepoint on successful and unsuccessful GS trials (collapsed across Side). Excitability is higher across time points on unsuccessful GS trials (*P* < 0.001). Stop signal presented 250 ms before target. Values are means ± standard error.

Median MEP data mirrored the same pattern of results with a main effect of Success (*F*_1,12_ = 15.693, *P* = 0.002, □_p_^2^ = 0.567, □_G_^2^ = 0.06), a main effect of Stimulation Time (*F*_6,72_ = 4.788, *P* = 0.025, □_p_^2^ = 0.285, □_G_^2^ = 0.09), and a Success × Stimulation Time interaction trending towards significance (*F*_6,72_ = 2.213, *P* = 0.096, □_p_^2^ = 0.156, □_G_^2^ = 0.02). In the median data, the Success × Side × Stimulation Time interaction also approached significance (*F*_6,72_ = 1.953, *P* = 0.084, □_p_^2^ = 0.140, □_G_^2^ = 0.01) but there were no other main effects or interactions (all P > 0.099).

It is possible that overall higher baseline EMG across unsuccessful compared to successful GS trials may have contributed to the difference between conditions in the corresponding MEP data. There was a main effect of Success (*F*_1,12_ = 14.295, *P* = 0.003, □_p_^2^ = 0.544, □_G_^2^ = 0.04) with baseline EMG on unsuccessful trials (2.9 ± 0.2 μV) higher than successful GS trials (2.3 ± 0.3 μV, t_12_ = −3.789, *P* = 0.003). There was a main effect of Stimulation Time (*F*_6,72_ = 5.756, *P* = 0.002, □_p_^2^ = 0.324, □_G_^2^ = 0.06) and the main effect of Side approached significance (*F*_1,12_ = 3.798, *P* = 0.075, □_p_^2^ = 0.240, □_G_^2^ = 0.04), but there were no significant interactions (all *P* > 0.128). When collapsed across Side and Success, baseline EMG increased from −150ms (2.0 ± 0.3 μV) at −25ms (3.1 ± 0.4 μV), which remained facilitated at 0ms (both *P* < 0.010).

## Discussion

We are the first study to use the dual-coil TMS technique to directly compare bilateral CME within the same trial of a response inhibition task. This has enabled us to reveal the CME dynamics of both response components simultaneously for the first time during partial cancellation of bimanual movement. MEP suppression was evident in equivalent amplitude and latency for both the responding and cancelled fingers during partial response inhibition. Concurrent suppression confirmed our first hypothesis and extended previous findings (Cowie et al., 2016; MacDonald et al., 2014). The subsequent divergence of CME between fingers confirmed our second hypothesis and supports a faciliatory input being selectively targeted to the unimanual response that is still required. The third hypothesis tested was that when a non-selective stopping mechanism cannot be adequately implemented and MEP suppression is not adequately applied in response to the stop cue, it is not possible to functionally uncouple the two fingers and change to selectively targeting just one side. This results in the default bimanual response being incorrectly executed. Our results support this hypothesis as CME was generally higher for both fingers during unsuccessful compared to successful partial cancellation. This result, along with faster responses on unsuccessful compared to successful partial cancellation trials, dovetails with the seminal horse-race model prediction (Logan & Cowan, 1984) that these trials were unsuccessful because faciliatory input for the bimanual response was too far progressed at the time of the stop cue. Although our results could not rule out the possibility that overall higher levels of background muscle activity caused the larger MEP amplitudes on unsuccessful trials. Overall, the results add substantial weight to behavioural (Bissett & Logan, 2014; Coxon et al., 2007; Coxon, Stinear, & Byblow, 2009), neurophysiological (Cowie et al., 2016; MacDonald et al., 2014) and computational modelling (MacDonald, McMorland, Stinear, Coxon, & Byblow, 2017) evidence of a non-selective stopping mechanism being engaged during partial response inhibition. Our results validate the use of dual-coil near-simultaneous TMS technique in the context of response inhibition. Furthermore we provide additional evidence that CME is unable to be inhibited selectively during the initial stages of partial cancellation of a bimanual movement, despite the presentation of a behaviourally selective stop cue.

The prepared bimanual response was cancelled via bilateral inhibition. When a selective stop cue was processed, a comparable decrease in CME occurred at the same time in both fingers, despite only the right being cued to stop. The equivalency between fingers within a single trial contributes strong evidence towards a non-selective stopping mechanism being engaged during partial response inhibition. However, in accordance with our third hypothesis, if CME was not reduced bilaterally to a sufficient degree a bimanual response was still made in error. We interpret the reduction in CME as evidence of the same neural inhibitory process being engaged (Aron & Poldrack, 2006; Boecker et al., 2011; Cai & Leung, 2011; Jahfari et al., 2011; Schmidt, Leventhal, Mallet, Chen, & Berke, 2013) and having a transient effect on motor output, before being overridden by the initial bilateral facilitatory drive. The transient inhibitory effect would explain the bimanual responses on unsuccessful GS trials being slower than bimanual responses on GG trials. Thus, there needs to be a substantial non-selective decrease in CME to some threshold (Badry et al., 2009) to cancel prepared bimanual actions. Such a threshold may be contextualised as a behavioural threshold for the stopping process (Frank, 2006) or an elevated level of neural inhibitory input that raises an activation threshold to surpass the initial bimanual facilitatory drive (MacDonald et al., 2017).

Facilitatory input was selectively targeted to the left finger motor representation to initiate a unimanual response. Following simultaneous suppression, CME for the left and right fingers diverged 25 ms prior to the target when CME for the responding left finger was higher. There are a few possible, non-exclusive reasons the divergence between fingers was less robust (i.e., significant in median but not mean MEP data) at the last stimulation timepoint (0 ms relative to target). It may be due to increased variance at this timepoint from inter-individual differences in movement re-initiation time. Alternatively, it may represent some level of cross-facilitation that is inherent to unimanual movement initiation (Muellbacher, Facchini, Boroojerdi, & Hallett, 2000). Finally, it may indicate that uncoupling of the two fingers had not been entirely achieved by this timepoint. Recent evidence does suggest uncoupling of fingers occurs later in the more difficult partial cancellation condition when only the non-dominant hand is responding (MacDonald, Laksanaphuk, Day, Byblow, & Jenkinson, 2021) such as in the current study. Future studies are needed to discriminate between these possibilities.

Evidence of non-selectively stopping a bimanual movement and selectively initiating a controlled unimanual response aligns with the stop-change/task switching framework (for a review see Boecker et al. 2013). Stop-change tasks involve beginning a prepared movement, stopping all prepared components of the movement, and then switching to a new alternate response via reprogramming. Evidence suggests these three processes are performed serially and function independently of each other, where the new alternate response is only activated after the stop process has terminated (Boecker, Gauggel, & Drueke, 2013; Verbruggen, McLaren, & Chambers, 2014; Verbruggen, Schneider, & Logan, 2008). Viewing the serial neural processes during partial response inhibition through the stop-change framework would explain the robust behavioural delay. Indeed, the neural networks implicated in the ‘stop’ part of the stop-change model are very similar to the non-selective prefrontal-basal ganglia pathway recruited for simple response inhibition i.e. the hyperdirect pathway (Aron, Herz, Brown, Forstmann, & Zaghloul, 2016; Boecker et al., 2011; Coxon et al., 2009; Elchlepp & Verbruggen, 2017; Frank, 2006; Isoda & Hikosaka, 2007; Kenner et al., 2010; Rangel-Gomez, Knight, & Kramer, 2015; Wiecki & Frank, 2013). Mounting evidence suggests the pre-supplementary motor area (preSMA) is important for resolving conflict in the presence of competing response options and reprogramming responses during task switching (Dove, Pollmann, Schubert, Wiggins, & von Cramon, 2000; Isoda & Hikosaka, 2007; Neubert, Mars, Buch, Olivier, & Rushworth, 2010). The preSMA therefore plays a pivotal role in the stop-change framework (Antons et al., 2019; Jha et al., 2015; Roberts & Husain, 2015; Swann et al., 2012). Of note, preSMA activation is specifically linked to successful partial cancellation trials of the ARI task (Coxon et al., 2016; Coxon et al., 2009; Coxon, Van Impe, Wenderoth, & Swinnen, 2012) which implies the presence of response reprogramming. Partial response inhibition behaviour can therefore be viewed at a neural level as a sequential non-selective stop, change/switch, then go process.

There is one key aspect in which partial response inhibition deviates from the traditional stop-change framework. Stop-change tasks do not require participants to change from bimanual to unimanual responses i.e. the alternate response is not a subset of the original response. For example, Boecker et al. (2011) used a stop-change task that required participants to inhibit the movement of their index and ring fingers and instead initiate a new movement with the middle finger. The new middle finger response was considerably faster than the default index and ring finger responses. This is in contrast to the robust delay from the responding finger in successful GS compared to GG trials of the ARI task (Cowie et al., 2016; Coxon et al., 2016; Coxon et al., 2007; MacDonald et al., 2014; Macdonald, Stinear, & Byblow, 2012; Wadsley et al., 2019). We posit that this behavioural deviation from the stop-change framework is due to the additional step of uncoupling the bimanual response to allow selection and re-initiation of a subset of the movement. In this way, partial response inhibition may be viewed as an extension of the stop-change framework that also incorporates bimanual coupling.

The study findings should be interpreted in light of some limitations. Firstly, the wide range of stimulation timepoints needed to record the dynamic CME curve combined with success rates around 50%, resulted in a somewhat restricted number of trimmed MEPs for averaging on successful and unsuccessful GS trials. Unfortunately, within practical time constraints we could not substantially increase trial numbers. Therefore, it was important to confirm all results with median MEP data, and in this way we have ensured robust and valid findings from the trimmed mean MEPs. It is also worth noting that our CME curves based on trimmed mean MEPs replicated those from previous studies (Cowie et al., 2016; MacDonald et al., 2014), adding further validity to the current findings. However, in future studies it may be prudent to focus on the most important time points (e.g. −100 −75 −25 0 ms relative to target) to increase the number of MEPs collected at each data point. Secondly, some participants were excluded from the analysis investigating the effect of stopping success as they were unable to provide a complete data set for unsuccessful GS trials. The reduced participant number (N = 13) led to this aspect of the analysis being underpowered (minimum of N = 15 required for power of 0.8). In future, to specifically investigate the neural processes that make a partial cancellation trial unsuccessful, the stop time could be set closer to the target to increase the proportion of unsuccessful trials when a bimanual response is still observed.

### Conclusion

The novelty of the current findings arises from using a dual-coil TMS technique to simultaneously record - and therefore directly compare - CME in the cancelled and responding hands within the same trial during a response inhibition task. Our results provide strong evidence that inhibition is unable to be applied selectively at a neural level during partial cancellation of a bimanual movement, despite the presentation of a behaviourally selective stop cue. Neural modulation during partial response inhibition can instead be viewed in light of the stop-change framework as a sequential non-selective stop, uncouple and switch, then selectively go process. The dual-coil TMS technique may prove especially useful to investigate these response inhibition mechanisms in patients, as it doubles the amount of data collected per trial compared to single-coil techniques and may therefore decrease session duration to something more tolerable in patient populations.

## Notes

*Conflict of interest:* The authors declare no competing financial interests.

### Competing Interest Statement

The authors have declared no competing interest.

